# A general model for the seasonal to decadal dynamics of leaf area

**DOI:** 10.1101/2024.10.23.619947

**Authors:** Boya Zhou, Wenjia Cai, Ziqi Zhu, Han Wang, Sandy P. Harrison, I. Colin Prentice

**Author notes:** Correspondence to*: Boya Zhou.

## Abstract

Leaf phenology, represented at ecosystem scale by the seasonal dynamics of leaf area index (LAI), is a key control on the exchanges of CO_2_, energy and water between the land and atmosphere. Robust simulation of leaf phenology is thus important for both dynamic global vegetation models (DGVMs) and land-surface representations in climate and Earth System models. There is no general agreement on how leaf phenology should be modelled. However, a recent theoretical advance posits a universal relationship between the time course of “steady-state” gross primary production (GPP) and LAI – that is, the mutually consistent LAI and GPP that would pertain if weather conditions were held constant. This theory embodies the concept that leaves should be displayed when their presence is most beneficial to plants, combined with the reciprocal relationship of LAI and GPP via (a) the Beer’s law dependence of GPP on LAI, and (b) the requirement for GPP to support the allocation of carbon to leaves. Here we develop a global prognostic leaf phenology model, combining this theoretical approach with a parameter-sparse terrestrial GPP model (the P model) that achieves good fit to GPP derived from flux towers in all biomes, and a scheme based on the P model that predicts seasonal maximum LAI as the lesser of an energy-limited rate (maximizing GPP) and a water-limited rate (maximizing the use of available precipitation). The exponential moving average method is used to represent the time lag between leaf allocation and modelled steady-state LAI. The model captures satellite-derived LAI dynamics across biomes at both site and global levels. Since this model outperforms the 15 DGVMs used in the TRENDY project, it could provide a basis for improved representation of leaf-area dynamics in vegetation and climate models.

## 1 Introduction

Leaf phenology refers to the annual life cycle of leaves, including budburst and unfolding, growth, senescence and abscission (Xie et al., 2018). It has far-reaching consequences, constraining the geographic distributions of tree species (Chuine, 2020) and strongly influencing the global carbon and water cycles (Piao et al., 2019). Thus, leaf phenology is a key process for understanding and predicting the interactions between terrestrial ecosystems and climate (Zhao et al., 2013), and an essential element of land surface models.

The global controls of leaf phenology have gained increasing attention in recent years (Tang et al., 2016; Piao et al., 2019) but there is still no accepted way to represent phenology in global models. Most land surface models predict seasonal cycles of LAI as outcomes of carbon allocation. For example, the Community Land Model Version 5 (CLM5.0) first estimates the carbon flux to leaves, taking account of the effects of climate-related variables including soil temperature and moisture, then converts leaf carbon mass to the equivalent LAI assuming a fixed Specific Leaf Area (SLA) for each vegetation type (Lawrence et al., 2019). The Community Atmosphere Biosphere Land Exchange (CABLE) model (Xia et al., 2017) and the Canadian Terrestrial Ecosystem Model (CTEM) (Arora and Boer, 2005) divide vegetation growth into phenophases and prescribe different proportions of photosynthate to be allocated to leaves in each phase. Leaf area is then derived based on empirical relationships with biophysical variables, including SLA. In most land surface models, biophysical variables are set to fixed values for each vegetation type (Kato et al., 2013; Meiyappan et al., 2015; Vuichard et al., 2017; Walker et al., 2017; Lienert and Joos, 2018). This approach requires many empirical values to be specified, introducing considerable uncertainty for global-scale simulations (Xia, et al., 2017). Some other models, such as the Integrated BIosphere Simulator (IBIS) model (Yuan et al., 2014) and the Interactions between Soil, Biosphere, and Atmosphere (ISBA) model (Gibelin et al., 2006; Delire et al., 2020), model SLA more dynamically, allowing it to be modified by leaf nitrogen concentration and atmospheric carbon dioxide. However, all of these approaches are *ad hoc*, lacking an explicit theoretical basis; and indeed, theory for the seasonal dynamics of carbon allocation is still incomplete (Franklin et al., 2012; Hartmann et al., 2020) and thus unable to provide a well-founded basis for modelling phenology.

An extensive literature deals with the timing of phenological transitions, most commonly the start of season (SOS) and end of season (EOS), and many models have been published describing the controls of these transitions. Current models for spring phenology in cold-winter climates are of two main types: one-phase models only consider the need for plants to accumulate heat during the ecodormancy phase, when bud regrowth is hampered by unfavourable external conditions (Reaumur, 1735; McMaster and Wilhelm, 1997; Zhou et al., 2018); two-phase models also consider the amount of chilling during the (preceding) endodormancy phase, which is an internally regulated process, whereby growth is suppressed even in favourable environmental conditions (Cannell and Smith, 1983; Chuine et al., 2000; Caffarra et al., 2011; Fu et al., 2020). Some other models include the influence of environmental factors on more complex aspects of phenology, such as the DormPhot model (Caffarra et al., 2011) which considers the effects of chilling, forcing, and photoperiod on dormancy induction and phase of forcing simultaneously. However, these phenology models are largely species- or plant type-specific and therefore not well suited to application in a global modelling context.

Experiments have shown that air temperature is the main driver of phenological changes in northern temperate and high-latitude regions (Fu et al., 2019; Meng et al., 2021), while precipitation is the major driver in drylands (Currier and Sala, 2022) and has a major impact on their spring phenology (Castillioni et al., 2022). The cyclical patterns of solar radiation and vapour pressure deficit are the two most important environmental variables linked to the production of new leaves and the shedding of old leaves in tropical evergreen forests (Chen et al., 2019). For robust global-scale modelling, it would be useful to seek an approach that could account for such dependencies via a single mechanism applicable to all biomes. In this spirit, Jolly et al. (2005) proposed the Growing Season Index Model, which considers photoperiod, vapour pressure deficit and air temperature as predictors. But even though this model – and others derived from it (Schaphoff et al., 2018) – have had some success in simulating seasonal leaf area dynamics, they are still limited in climate-change applications because of their lack of theoretical underpinnings.

Xin et al. (2018) proposed a promising alternative method for simulating the seasonal dynamics of LAI, in which LAI time series are estimated from the seasonal time course of GPP via the Beer’s law dependence of GPP on LAI (Swinehart, 1962) and the reciprocal requirement for GPP to support LAI development. The underlying principle – that seasonal variations in LAI are coordinated with variations in GPP – can be considered as an eco-evolutionary optimality hypothesis (Harrison et al., 2021), because it implies that leaves are displayed at (or near) the time when they are able to be most productive. A “semi-prognostic” LAI dynamics model (Xin et al., 2020) based on this approach was developed. It requires annual peak LAI data from satellite data as input, to provide an upper bound for predicted LAI. This model has been shown to capture both the spatial pattern and seasonal variation of satellite-derived LAI on a global scale, albeit with some limitations in high-latitude and tropical areas. Zhou et al. (2023) further developed the model of Xin et al. (2020) by adding a scheme to predict seasonal maximum LAI, thus allowing a fully prognostic (i.e. independent of satellite data) simulation of the seasonal time course of LAI. This model can generally capture the seasonal variation of LAI, although it performs less well for biomes with typically high LAI such as evergreen broadleaf forests and closed shrublands, and still requires nine parameters to be estimated separately for each biome.

Here we adopt the theoretical approach proposed by Xin et al. (2018, 2020) but in addition we build on the development of a universal, first principles-based model for GPP (the P model: Prentice et al., 2014; Wang et al., 2017; Stocker et al., 2020; Mengoli et al., 2022). The P model rests on eco-evolutionary optimality hypotheses that have been tested one-by-one using independent data. When forced by satellite-derived LAI data it achieves good representations of the seasonal cycle of GPP across all biomes (Stocker et al., 2020), and a realistic simulation of site-based GPP trends (Cai and Prentice, 2020), without requiring biome-specific parameters. We also apply a top-down algorithm for predicting seasonal maximum LAI (Zhu et al., 2023; Cai et al., 2024), which makes use of the P model. This algorithm is based on the principle that this quantity is limited either by available energy (in which case a profit-maximizing criterion applies), or by water availability (when it is limited by the transpiration demands of photosynthesis). This allows us to build a fully prognostic solution for simulating LAI time series, which combines the robustness of the P model with our top-down criteria for seasonal maximum LAI and the conceptual simplicity and generality of the approach pioneered by Xin et al. (2018, 2020).

The present study has two objectives: (1) to develop a global leaf phenology model, independent of satellite data, that can simulate seasonal to multidecadal dynamics of LAI both at eddy-covariance flux sites and globally; and (2) to evaluate the model across biomes, using both *in situ* and satellite measurements.

## 2 Data and methods

### 2.1 Model overview

The modelling workflow has two steps (Fig. 1): (1) Derivation of steady-state LAI dynamics from the time course of potential GPP through the Beer’s law dependence of GPP on LAI, and the requirement for GPP to support LAI development. (2) Simulation of actual LAI dynamics, allowing for the time lag of leaf allocation behind steady-state LAI.

**Figure 1.**
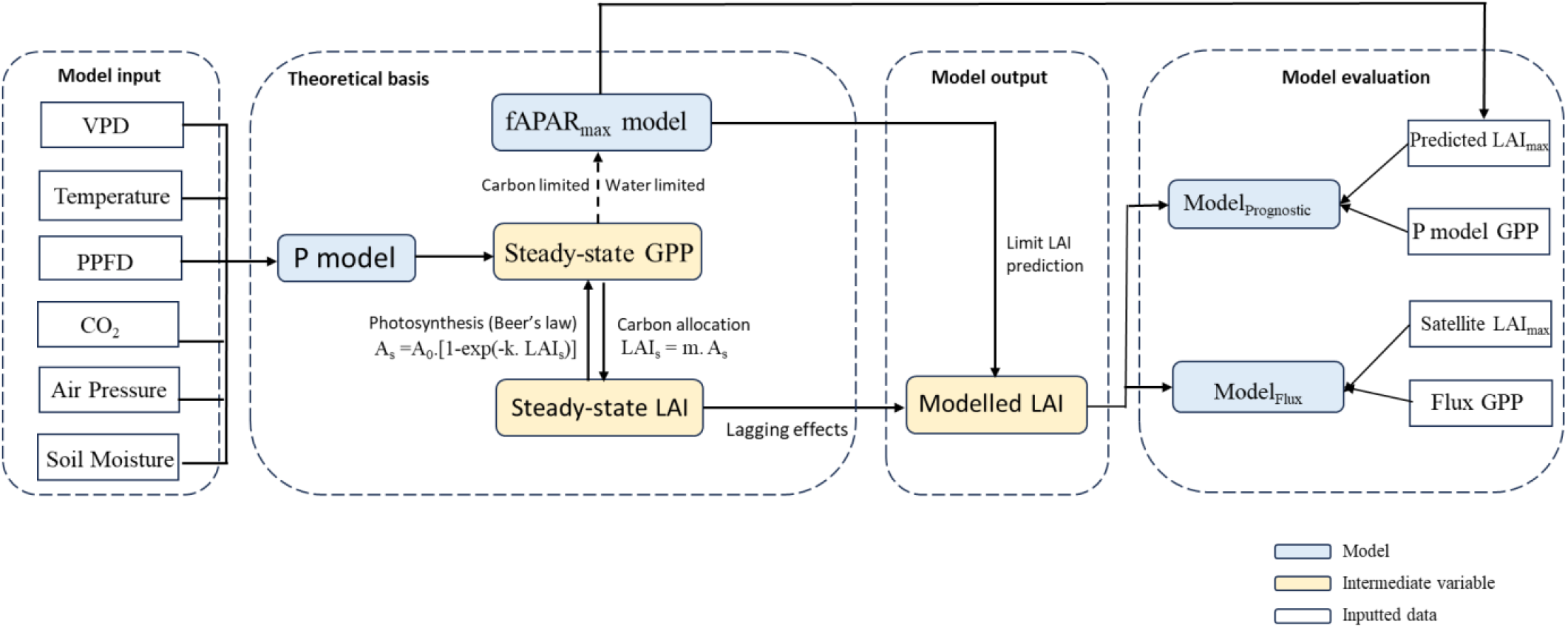
Workflow of the LAI model and its evaluation. Notation: *A*_*s*_, steady-state GPP (g C m^−2^ day^−1^); *A*_0_, potential GPP (g C m^−2^ day^−1^); LAI_s_, steady-state LAI (m^−2^ m^−2^); LAI_max_, annual peak LAI (m^2^ m^−2^). *A*_s_ and LAI_s_ are the mutually consistent GPP and LAI that would be supported if environmental conditions were held constant. *A*_0_ is the GPP that would be supported under the same conditions if all light incident on the canopy were absorbed by leaves, i.e. as LAI −> ∞. *k* is the light extinction coefficient (≈ 0.5) and *m* is a quantity (to be determined) relating *A*_s_ and LAI_s_.

#### 2.1.1 Prediction of annual maximum LAI

We treat fAPAR (the fraction of incident photosynthetically active radiation that is absorbed by leaves) and LAI as interchangeable, assuming they are monotonically related by Beer’s law (Swinehart, 1962; Fig. 1). Global spatial patterns and temporal trends of the seasonal maximum value of fAPAR (fAPAR_max_) or LAI (LAI_max_) can be predicted to first order based on eco-evolutionary optimality principles (Zhu et al., 2023; Cai et al., 2024). fAPAR_max_ can be represented as the lesser of two quantities. One comes from the energy-limited equation, which assumes that there is an optimal LAI at which net carbon profit is maximized after accounting for the costs of building and maintaining leaves and the total below-ground allocation required to support those leaves. The other comes from the water-limited equation, which assumes that vegetation cover can be limited by precipitation because of the tight coupling between transpiration and photosynthesis. The costs in the energy-limited equation are not explicitly modelled; instead, they are subsumed in a single constant (*z*), whose value has been estimated from satellite-derived LAI data. Similarly, the water-limited equation contains a single parameter (*f*_0_), representing the fraction of annual precipitation that can be transpired by plants. In the original fAPAR_max_ framework (Zhu et al., 2023; Cai et al., 2023) *f*_0_ was assigned a single constant value, also estimated from satellite LAI data, equal to 0.62. Here, we have substituted this constant value with an empirical equation that relates *f*_*0*_ to the climatological Aridity Index (AI) (Good et al., 2017), thus allowing *f*_0_ to decline as AI increases beyond a value close to the separation between energy- and water-limited regimes. The optimal fAPAR can then be represented by Eqn 1 and the optimal LAI by Eqn 2:

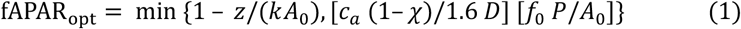

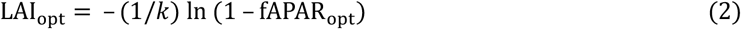

In Eqn 1, the first part represents the energy-limited equation, the second part the water-limited equation. *k* is the light extinction coefficient (set at 0.5), and *A*_*0*_ is the potential GPP i.e. the GPP that would be achieved if fAPAR = 1. *c*_*a*_ is the ambient CO_2_ partial pressure (Pa), and *χ* is the ratio of leaf-internal CO_2_ partial pressure (Pa) to *c*_*a*_. *D* is the vapour pressure deficit (VPD, Pa) during the growing season and *P* is the annual total precipitation (mol m^−2^ year^−1^). Further information on model derivation, including the estimation of *z* and *f*_*0*_, is provided in Supplementary Information Note 1.

#### 2.1.2 Simulating the dynamics of LAI

GPP is dependent on LAI (with the relationship governed by meteorological conditions) since leaves are the main organs of photosynthesis. On the other hand, GPP represents the total rate of carbon fixation, of which some fraction accrues to leaves. A general principle governing leaf phenology is that it should allow plants to achieve competitive success through maximizing photosynthesis subject to environmental constraints (Franklin et al., 2012). Based on the mutual relationship between GPP and LAI, Xin et al. (2018) proposed a criterion consistent with this principle, based on the concepts of steady-state LAI and GPP − that is, the LAI and GPP that would be in equilibrium if weather conditions were constant (Eqn 3). Given daily meteorological conditions, the steady-state LAI is modelled by solving the closed system of equations as follows (Xin et al., 2020) (Eqn 3 and Eqn 4):

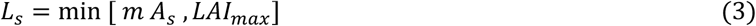

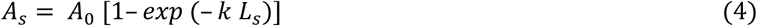

where *L*_s_ denotes steady-state LAI; *m* denotes the fraction of GPP allocated to LAI; and *A*_s_ denotes the steady-state GPP. The solution is given by Eqn 5:

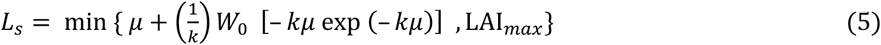

where *μ* = *mA*_0_ and *W*_0_ is the principal branch of the Lambert W function. Further information on the solution of Eqns 3 and 4 is provided in Supplementary Information Note 2.

The processes governing canopy photosynthesis and vegetation phenology do not respond instantaneously to weather fluctuations, so there are inherent delays between the steady-state LAI and the real-time dynamic LAI. Although photosynthesis responds within minutes, the allocation of photosynthate to leaves and other tissues can take from days to several months (Mengoli et al., 2022; Sierra et al., 2022). Thus, it is reasonable to model GPP on an hourly to daily basis, and to simulate leaf-area dynamics as a slower process, lagging steady-state GPP. Without such a lag, LAI would fluctuate unrealistically from day to day. We adopt the exponential weighted moving average method (Eqn 6) to incorporate this lag effect (Mengoli et al., 2022). This method is widely used in models to smooth out short-term fluctuations and highlight longer-term trends or cycles. It assigns exponentially decreasing weights to older LAI values, making it more responsive to recent changes in the data compared to a simple moving average (Yu et al., 2020):

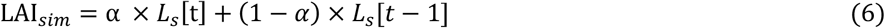

where the simulated actual LAI (LAI_sim_) is a weighted average of the steady-state LAI value corresponding to the steady-state GPP of the day (*L*_s_) and its acclimated value from the previous day, with weightings of α and (1 − α) respectively. α is set to 0.067 here, corresponding to approximately 15 days of memory. We also tested a range of alternative values of α (0.33, 0.143, 0.1, 0.067, 0.05, 0.04, 0.033, 0.022 and 0.0167, corresponding to 3, 7, 10, 15, 20, 25, 30, 45 and 60 days) (Table S2; Fig. S1). Further information on the derivation of Eqn 6 and the parameter α are provided in Supplementary Information Note 3.

#### 2.1.3 Estimation of *m*

Because actual LAI and GPP lag behind *L*_*s*_ and *A*_*s*_, the parameter *m* cannot be estimated directly from daily or monthly data (Xin et al., 2016; Xin et al., 2019). However, a natural approximation for *m* is the ratio of annual average LAI to annual mean GPP:

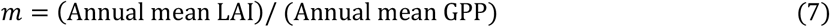

Xia et al. (2015) found that annual total GPP can be robustly estimated as a constant fraction (around 2/3) of the product of the length of the CO_2_ uptake period (growing season length, GSL) and the seasonal maximum of GPP (GPP_max_). Applying the same approach to LAI, we estimate *m* as follows:

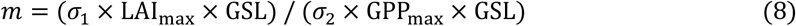

where LAI_max_ is the annual maximum LAI and *σ*_1_ and *σ*_2_ are parameters. We further assume that GPP_max_ is achieved when both *A*_0_ and fAPAR achieve their peak values, that is GPP_max_ = *A*_0max_ × fAPAR_max_, and that *A*_0max_ also follows the Xia et al. (2015) theory and can be represented as *A*_0ma*x*_ = *A*_0sum_ / (σ_3_ × GSL), where *A*_0sum_ is the annual sum of *A*_0_. Thus:

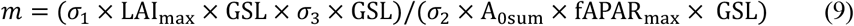

Combining parameters (defining *σ* = *σ*_1_ × *σ*_3_ / *σ*_2_), we obtain:

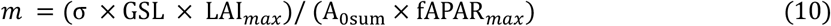

We estimate GSL as the length of the continuous period with mean daily temperatures > 0°. LAI_max_ is the seasonal maximum LAI, from Eqn 2; *A*_0sum_ is the annual total potential GPP; fAPAR_max_ is the seasonal maximum fAPAR, from Eqn 1; and *σ* represents the extent to which seasonal LAI dynamics depart from a “square wave” whereby maximum LAI would be maintained over the whole growing season (Xia et al., 2015; Zhou et al., 2023). To estimate *σ* we used data at flux sites to obtain “observed” values of *m*, as the ratio of annual average (satellite-derived) LAI to annual mean (flux tower-derived) GPP, and “observed” total *A*_0_, which is the ratio of flux tower-derived GPP to satellite-based fAPAR (data sources are described below). Then *σ* was calculated according to Eqn 10 with “observed” *m* values, satellite-derived seasonal maximum LAI and fAPAR, and “observed” *A*_0_ as inputs. This method yielded *σ* = 0.771 as a global estimate.

### 2.2 Data and model evaluation

#### 2.2.1 Flux-tower data and site-based analysis

GPP and meteorological data at flux-tower sites were obtained from the FLUXNET 2015 Tier Two and ONEFLUX datasets (http://fluxnet.fluxdata.org/) (Pastorello et al., 2020). At site level, the sub-daily P model (Mengoli et al., 2022) was used to calculate potential GPP with the FLUXNET datasets providing the meteorological variables required as input: incident photosynthetic photon flux density (PPFD_IN, µmol m^−2^ s^−1^), vapour pressure deficit (VPD_F, Pa), air temperature (TA_F,° C), and carbon dioxide mole fraction (CO_2__F_MDS, µmol mol^−1^) on a half-hourly timestep. In contrast with the standard P model (Stocker et al., 2020), the sub-daily P model (Mengoli et al., 2022) implements acclimation of photosynthetic parameters to midday conditions, when the light is greatest, and explicitly separates the subdaily (fast) responses of photosynthesis and stomatal conductance from slower acclimated responses to environmental variations over a 15-day period. The effect of soil moisture stress on simulated GPP was considered separately by applying an additional empirical soil moisture limitation function (Mengoli et al., 2023).

Some days of climate data are missing from some sites due to quality control issues, so simulated GPP cannot be calculated for those days. Considering that the fAPAR_max_ model (Eqn 1) (Zhu et al., 2023; Cai et al., 2024) requires annual total potential GPP as input, only those site-years with >300 daily GPP simulations were used for analysis from a total of 149 flux tower sites. We used data in the time range from 2001 to 2018 (1038 site-years), including 47 evergreen needleleaf forest (ENF) sites, 14 evergreen broadleaf forest (EBF) site, 25 deciduous broadleaf forest (DBF) sites, seven mixed forest (MF) sites, two closed shrubland (CSH) sites, seven open shrubland (OSH) sites, six woody savanna (WSA) sites, six savanna (SAV) sites, and 31 grassland (GRA) sites (Table S1).

#### 2.2.2 Gridded climate data and global analysis

We used the standard P model (Stocker et al., 2020) for the global analysis. Daily precipitation, maximum, minimum and mean temperature, and surface downwelling shortwave radiation at 0.5° resolution, as inputs, were downloaded from the WFDE5 dataset version 2.1 (Cucchi et al., 2022). VPD was calculated using maximum and minimum temperature and water vapour pressure. Solar radiation was converted to incident PPFD assuming a flux: energy ratio of 4.6 μmol J^−1^ and a photosynthetically active fraction of 0.5. Atmospheric pressures were calculated from elevation using the barometric formula (Berberan-Santo et al., 2003). We downloaded globally averaged monthly mean CO_2_ concentrations (μmol mol^−1^) from NOAA Global Monitoring Laboratory for 2001-2019 (NOAA/GML; https://gml.noaa.gov/ccgg/trends/; last access November 2023) and then interpolated it into daily scale.

The effect of soil moisture stress on simulated GPP was represented using the empirical ‘penalty factor’ developed by Stocker et al. (2020), with the annual time course of soil moisture calculated using the Simple Process-led Algorithms for Simulating Habitats (SPLASH) model (version 1: Davis et al., 2017). As the standard P model makes separate calculations for GPP of plants following the C_3_ or C_4_ photosynthetic pathways, a dynamic C_4_ vegetation fraction was simulated based on a C_3_/C_4_ competition model (https://pyrealm.readthedocs.io/en/latest/) embedded in the P model. This model requires tree cover percentages as inputs, which were derived from MODIS MOD44B v006 during 2001-2019 (DiMiceli et al., 2015). Areas where cropland cover is >50% were excluded from the C_3_/C_4_ vegetation fraction map. Cropland cover data at 0.05° resolution from 2001-2019 were derived from MODIS MCD12C1 v006 (Friedl et al., 2015).

#### 2.2.3 Satellite LAI data (Model evaluation data)

We used LAI time series derived from two alternative datasets (MODIS and Copernicus LAI) to evaluate the model’s performance. MODIS LAI data were derived from the MODIS MOD15A2H Leaf Area Index/FPAR product, given at a resolution of 0.05° resolution at daily timestep from 2001−2019 (Jiang et al., 2016). Copernicus LAI data were derived from Copernicus Global Land Products (https://gbov.acri.fr) for the same period at 1 km resolution. The results described below are based on the MODIS data product. Results of model evaluation based on the Copernicus data are shown in the Supplementary materials (Fig. S2; Fig. S3).

#### 2.2.4 Simulated LAI data from the TRENDY project

We downloaded simulated LAI data during 2001−2019 from the ensemble of 15 TRENDY ecosystem models (TRENDY-v9; Table 1). The S2 simulations were used, in which identical, time-varying climate and CO_2_ are prescribed to all the models. The LAI datasets from different models were regridded to 0.5° resolution using the first-order conservative remapping function (remapcon) from the Climate Data Operators (CDO) software package (https://code.mpimet.mpg.de/projects/cdo).

**Table 1.** Details of the models from TRENDY-v9.

| Model name | Spatial resolution | Reference |
| --- | --- | --- |
| CABLE-POP | 1°×1° | Haverd et al. (2018) |
| CLASSIC | 2.8125°×2.8125° | Melton et al. (2020) |
| CLM 5.0 | 0.9375°×1.25° | Lawrence et al. (2019) |
| IBIS | 1°×1° | Yuan et al. (2014) |
| ISAM | 0.5°×0.5° | Meiyappan et al. (2015) |
| ISBA-CTrip | 1°×1° | Delire et al. (2020) |
| JSBACH | 1.875°×1.875° | Reick et al. (2021) |
| JULES-ES | 1.25°×1.875° | Wiltshire et al. (2021) |
| LPJ-GUESS | 0.5°×0.5° | Smith et al. (2014) |
| LPX-Bern | 0.5°×0.5° | Lienert and Joos (2018) |
| OCN | 1°×1° | Zaehle and Friend (2010) |
| ORCHIDEEv3 | 0.5°×0.5° | Vuichard et al. (2017) |
| SDGVM | 1°×1° | Walker et al. (2017) |
| VISIT | 0.5°×0.5° | Kato et al. (2013) |
| YIBs | 1°×1° | Yue and Unger (2015) |

#### 2.2.5 Evaluation methods

The LAI model depends on several components, including P model-derived GPP dynamics and the prediction of seasonal maximum fAPAR and LAI. We conducted multiple sets of simulations at flux sites to investigate the dependence of model performance on alternative model setups. Two model combinations were defined: (1) Model_Flux_: Inputs are flux-derived *A*_0_ and seasonal maximum LAI obtained from the remote sensing data. Here *A*_0_ is calculated by dividing flux-derived GPP by satellite-derived fAPAR (Table 1; Fig. 2b; Table 2; Fig. 3). (2) Model_Prognostic_: Inputs are P model-derived *A*_0_ and predicted seasonal maximum LAI from Eqn 2 (Table 1; Fig. 2a; Table 2; Fig. 3).

**Table 2.** Regression coefficient (Slope), *R*^2^ and RMSE of simulated and observed LAI assessed at different timescales. **Model**_**Prognostic**_: Simulated LAI dynamics using P model-derived GPP and simulated annual peak LAI from the fAPAR_max_ model as inputs. **Model**_**Flux**_, Simulated LAI dynamics using flux tower-derived GPP and annual peak satellite LAI as inputs.

|  | Model<sub>Prognostic</sub> |  |  | Model<sub>Flux</sub> |  |  |  |
| --- | --- | --- | --- | --- | --- | --- | --- |
| | Slope | $R^2$ | RMSE | Slope | $R^2$ | RMSE | N |
| 7 days | 0.88 | 0.60 | 0.76 | 0.80 | 0.67 | 0.76 | 53863 |
| Seasonal | 0.99 | 0.67 | 0.79 | 0.84 | 0.70 | 0.68 | 13184 |
| Annual | 0.96 | 0.70 | 0.61 | 0.90 | 0.87 | 0.29 | 1138 |
| Spatial | 0.98 | 0.72 | 0.47 | 0.93 | 0.90 | 0.26 | 163 |

**Figure 2.**
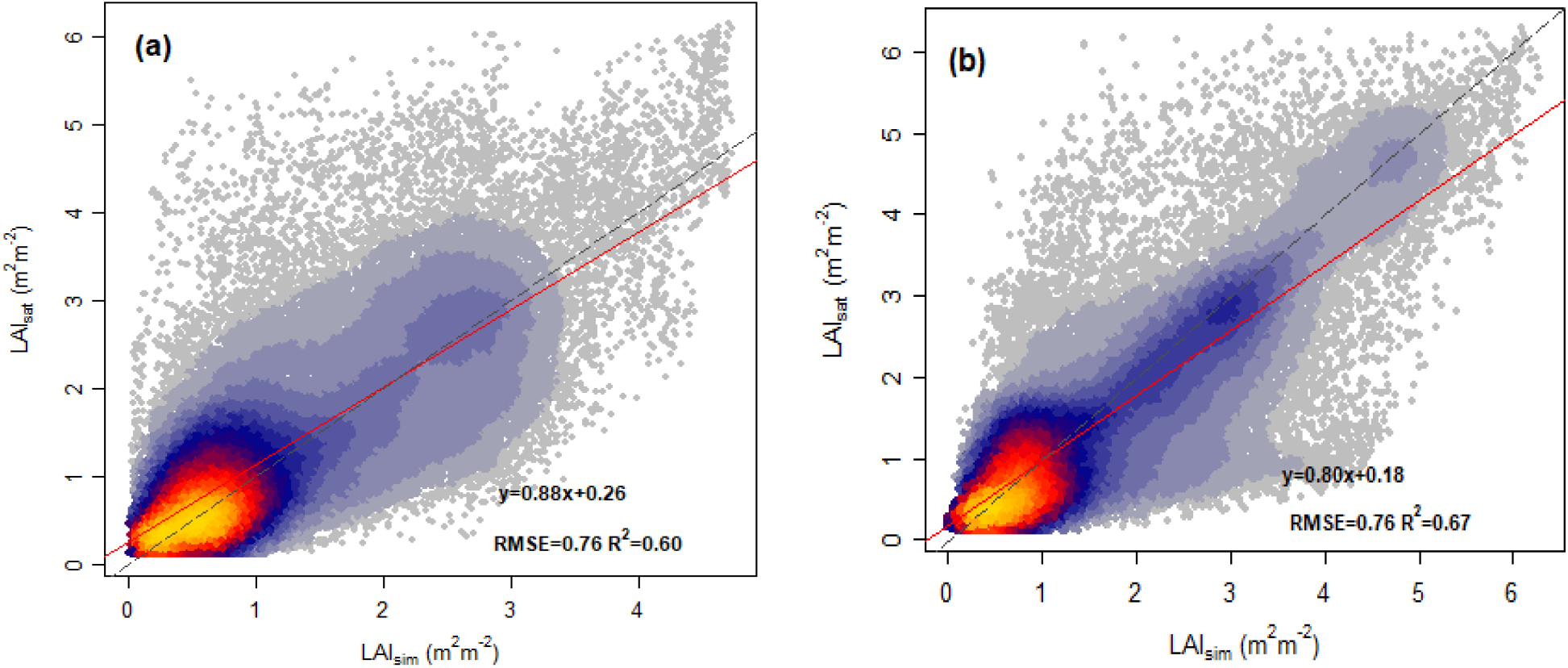
Correlation of observed and modelled seven-day mean LAI values for all sites pooled. **(a) Model**_**Prognostic**_: Simulated LAI dynamics using P model-derived GPP and simulated annual peak LAI from the fAPAR_max_ model as inputs. **(b) Model**_**Flux**_: Simulated LAI dynamics using flux tower-derived GPP and annual peak satellite LAI as inputs. Grey lines are the 1:1 line; red lines are the regression lines.

**Figure 3.**
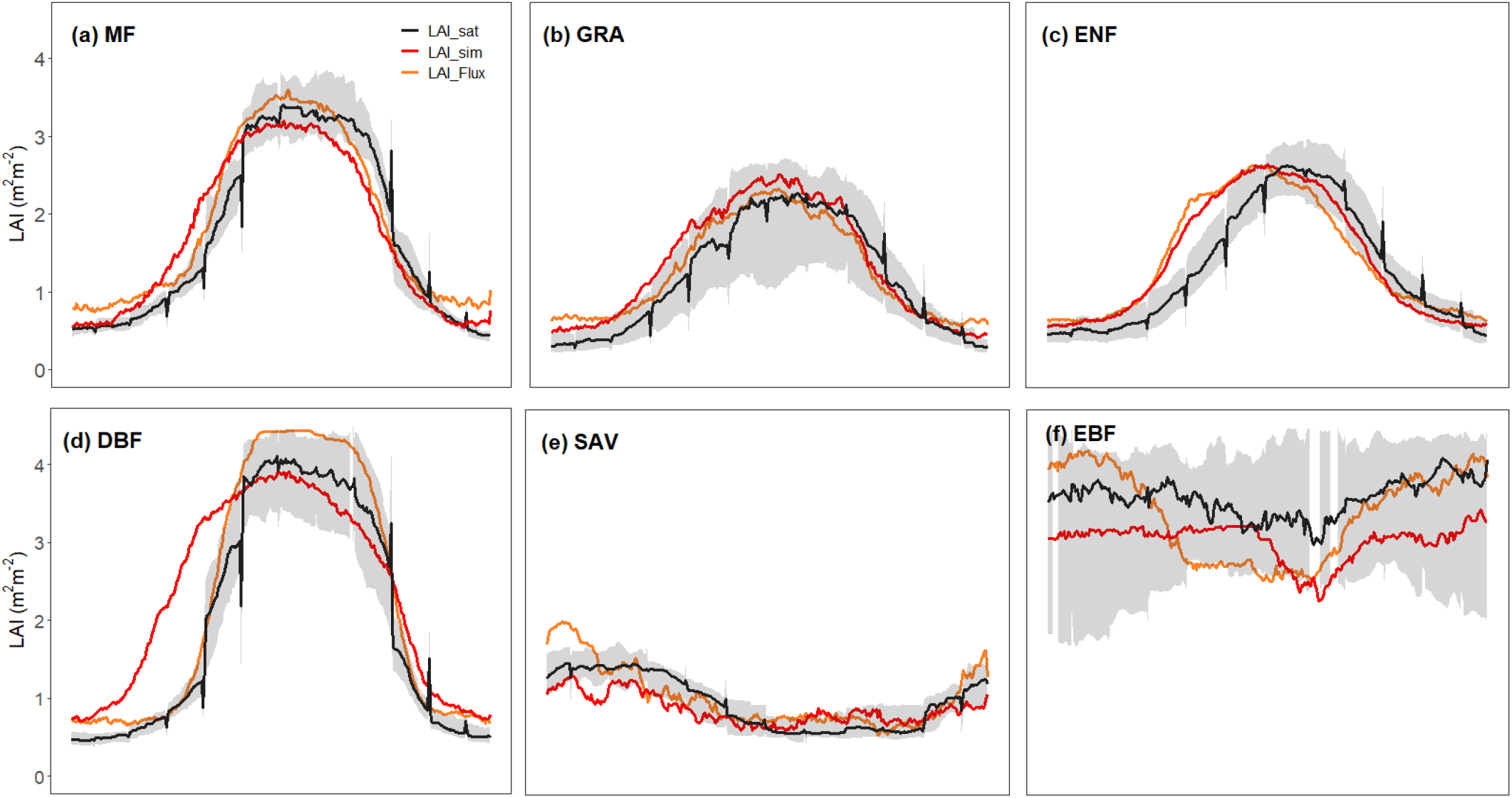
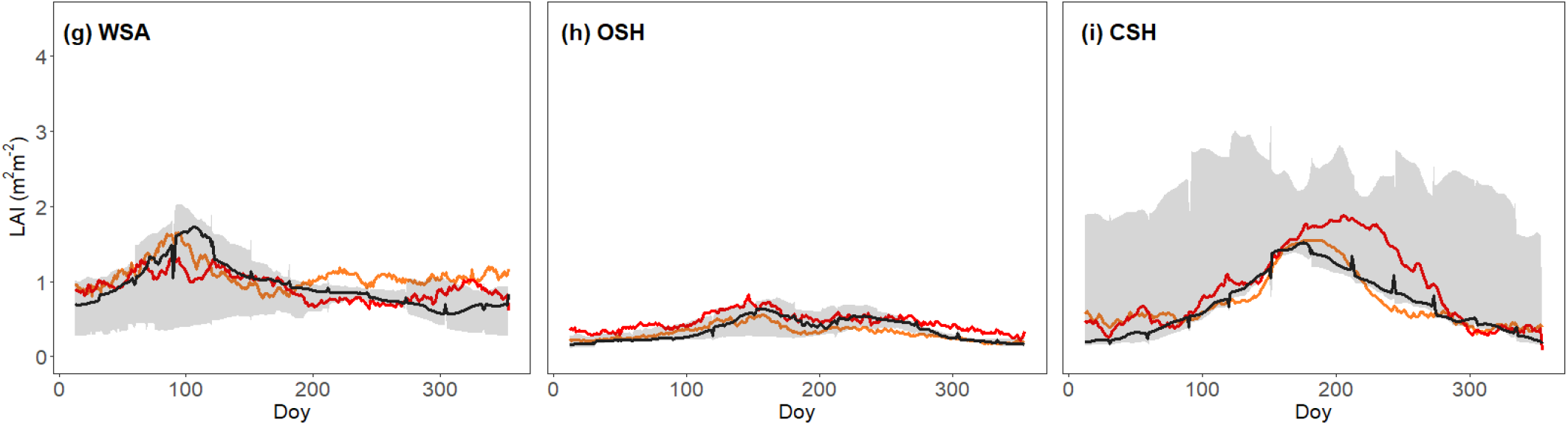
Mean seasonal cycles of LAI by biomes. Observations are shown by the black line and grey band, representing the median and 33/66 % quantiles of all data (multiple sites and years) pooled by biome. The red line (Model_Prognostic_) is the seasonal variation of simulated LAI across all sites and years for each biome, where simulated LAI is calculated using P model derived-GPP and annual peak LAI from the fAPAR_max_ model as inputs. The orange line (Model_Flux_) is the seasonal variation of simulated LAI across all sites and years for each biome, where simulated LAI is calculated using flux tower-derived GPP and annual peak MODIS LAI as inputs. **CSH**, Closed shrubland; **DBF**, Deciduous broadleaf; **EBF**, Evergreen broadleaf; **ENF**, Evergreen needleaf; **GRA**, Grassland; **MF**, Mixed Forest; **OSH**, Open shrubland; **SAV**, Savanna; **WSA**, Woody savanna.

We analysed spatial (multi-year mean values by site), annual, seasonal (mean by month of year), and weekly (mean by week of year) variability for both model combinations (Table 1), pooling data by biomes (Table S3). The models’ performance in simulating intra- and inter-annual LAI variability was also evaluated for different biomes (Fig. 3; Fig. 4). We separately analysed annual mean LAI (Fig. 5a, 5b) on an annual scale. We also evaluated the performance of our model and other 15 models participating in the TRENDY project in predicting multi-year average LAI (2001 to 2019), annual average LAI time series, and seasonal LAI variation by comparing with the LAI products derived from MODIS (Jiang et al., 2016).

**Figure 4.**
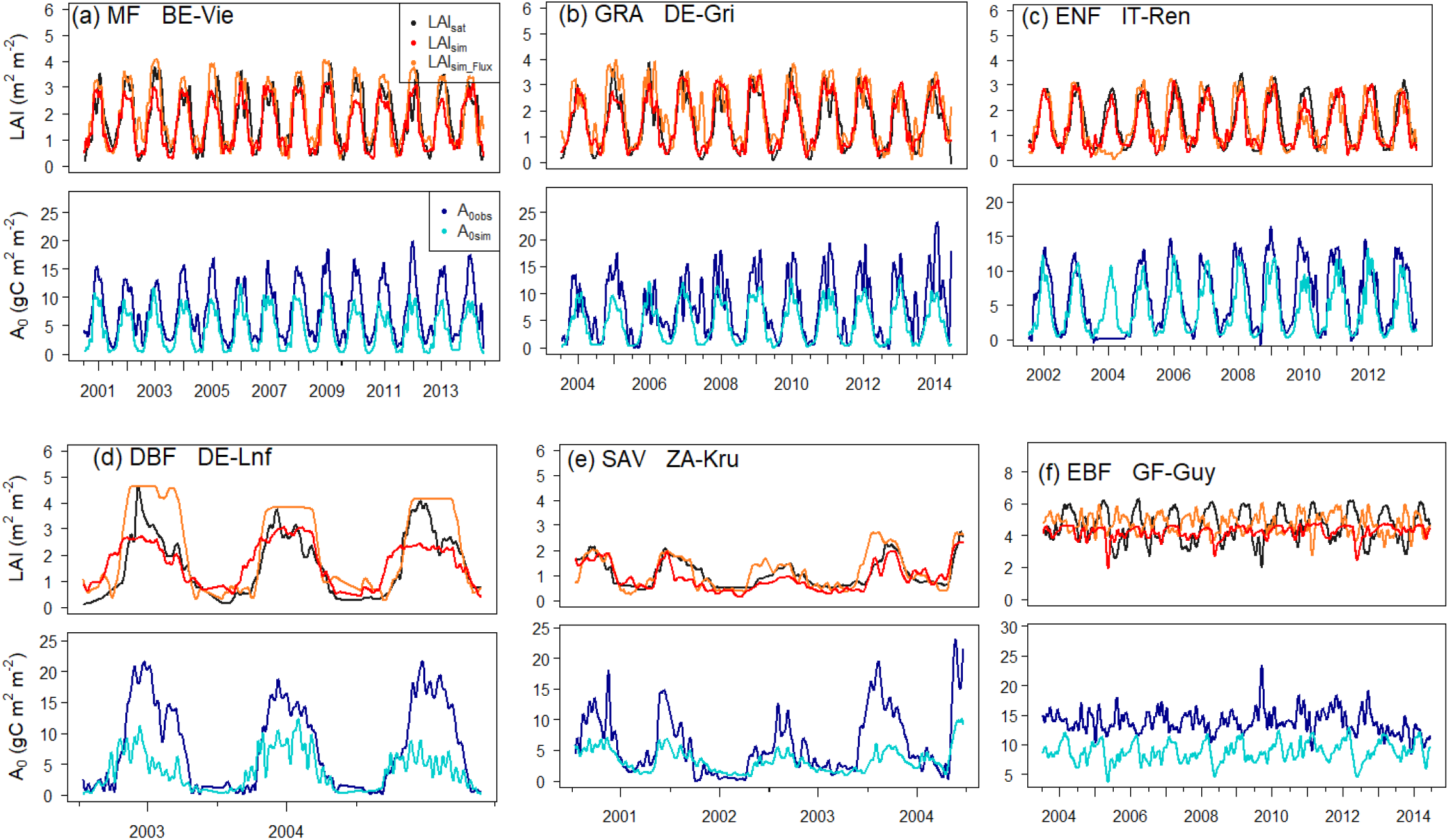
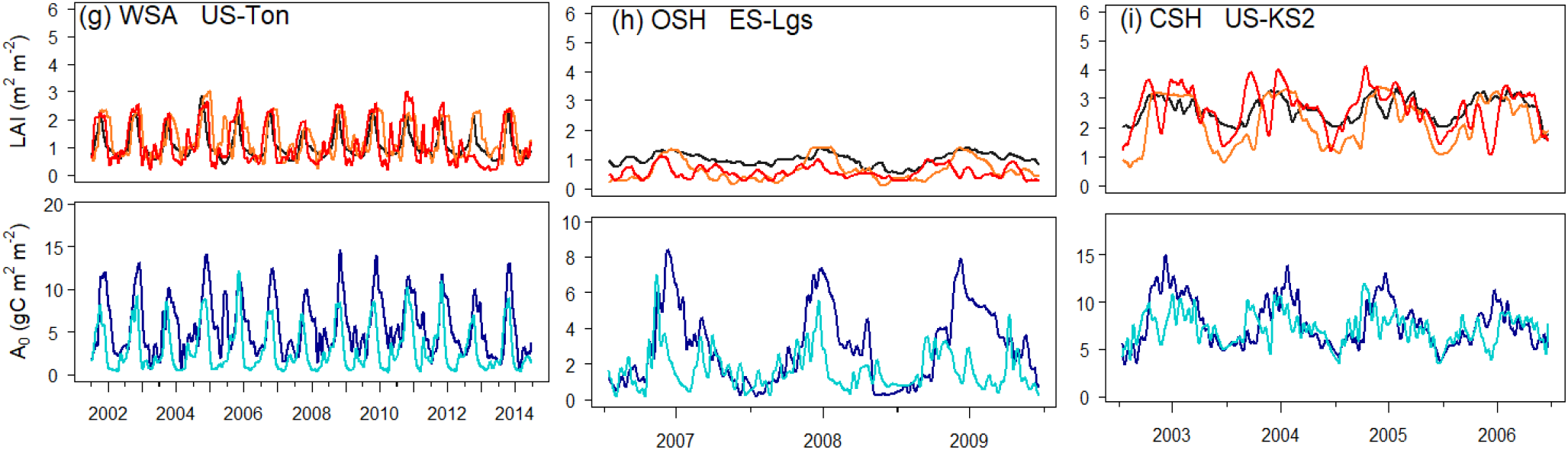
Representative time series of predicted and observed LAI and potential GPP in different biomes. LAI_sat_, Satellite LAI; LAI_sim_, Simulated LAI dynamics using P model-derived GPP and simulated annual peak LAI from the fAPAR_max_ model as inputs (Model_Prognostic_); LAI_sim_Flux_, Simulated LAI dynamics using flux tower-derived GPP and annual peak satellite LAI as inputs (Model_Flux_); A_0obs_: Observed potential GPP at flux sites (g C m^-2^ day^−1^); A_0sim_: Simulated potential GPP at flux sites (g C m^−2^ day^−1^).

**Figure 5.**
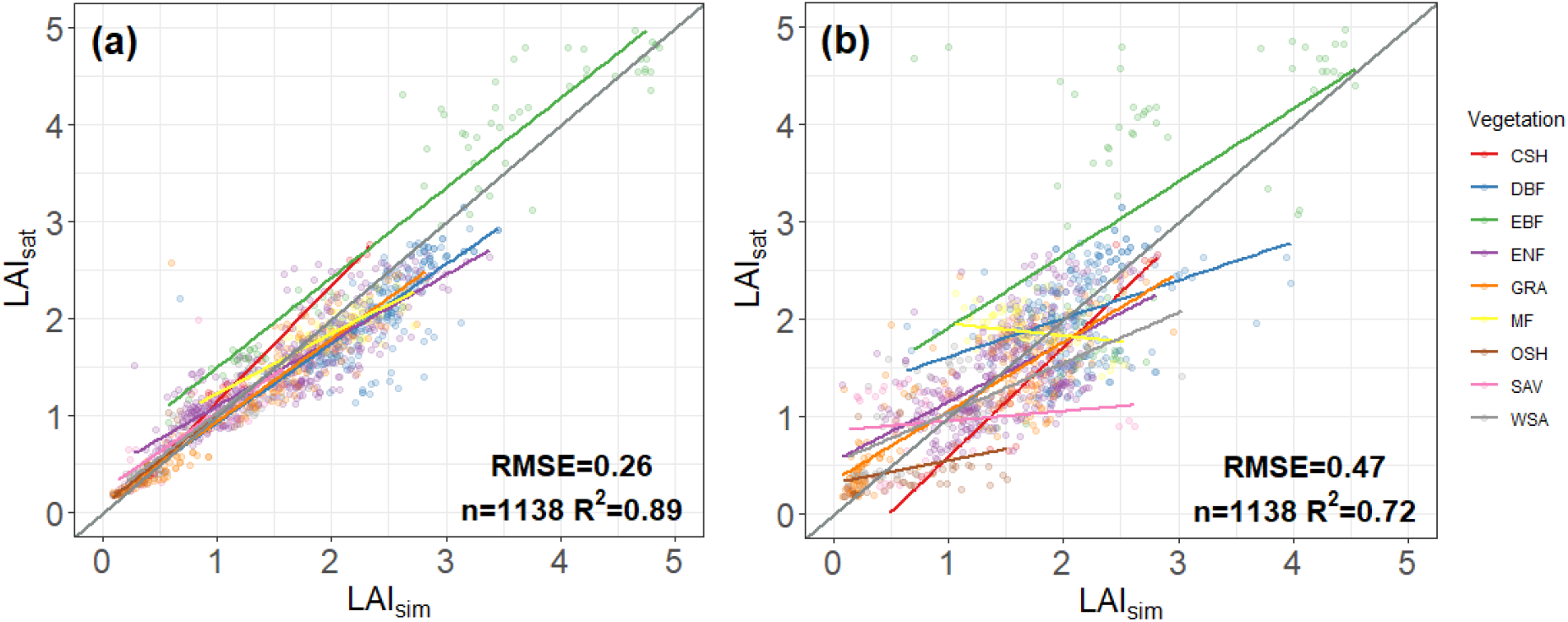
Regression analysis for **(a)** satellite-derived versus simulated annual average LAI, where simulated LAI is calculated using flux tower-derived GPP and annual peak MODIS LAI as inputs (Model_Flux_); **(b)** satellite-derived versus simulated annual average LAI, where simulated LAI is calculated using P model-derived GPP and annual peak LAI from the fAPAR_max_ model as inputs (Model_Prognostic_). Grey lines are the 1:1 lines and coloured lines are regressions basded on all sites within each biome. *R*^2^ and RMSE are based on pooled data for all biomes. **CSH**, Closed shrubland; **DBF**, Deciduous broadleaf; **EBF**, Evergreen broadleaf; **ENF**, Evergreen needleaf; **GRA**, Grassland; **MF**, Mixed Forest; **OSH**, Open shrubland; **SAV**, Savanna; **WSA**, Woody savanna.

**Figure 6.**
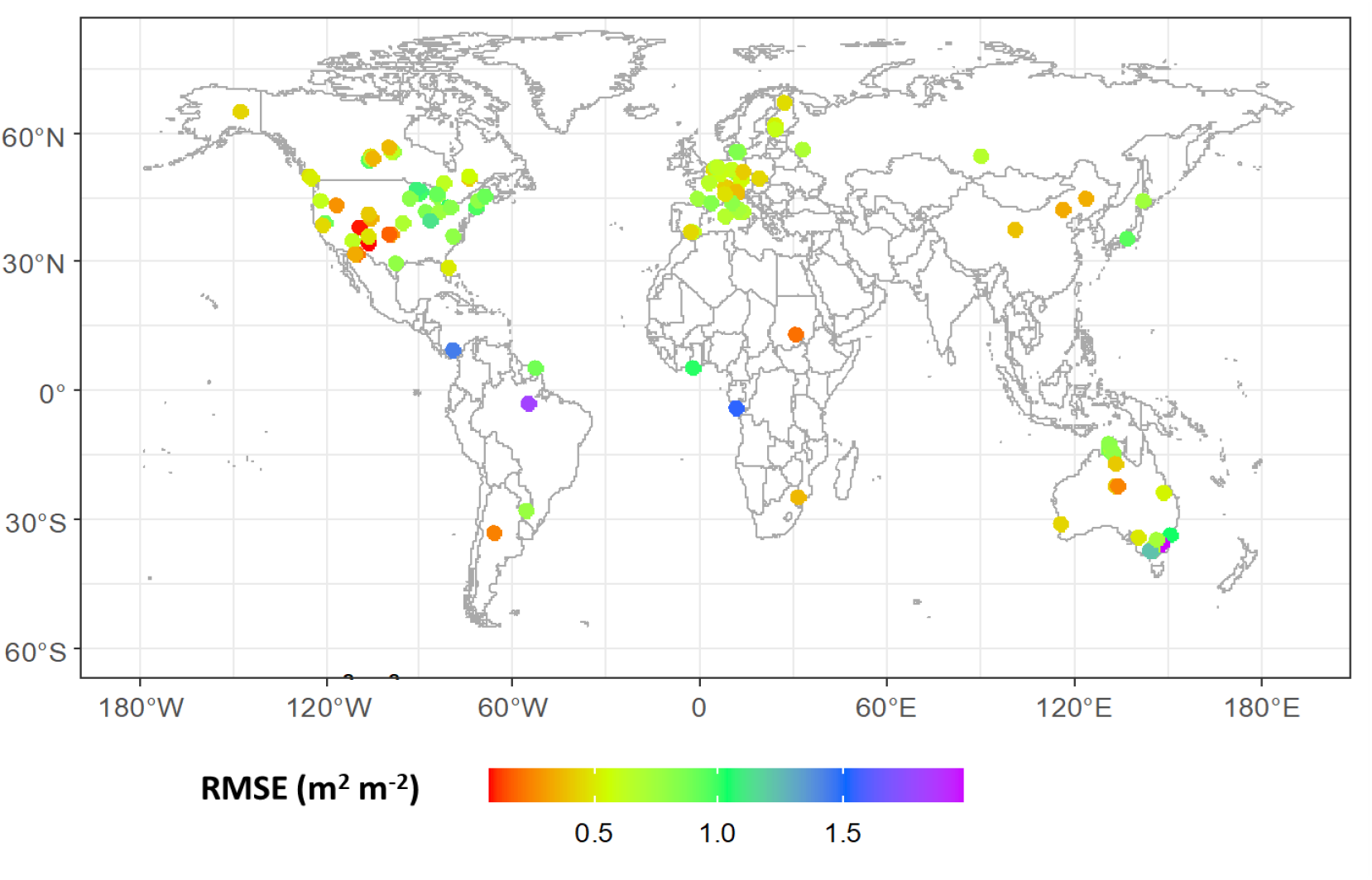
The spatial distribution of root mean square error (RMSE) between satellite-derived and modelled LAI from Model_Prognostic_ at each flux tower site.

We calculated *R*^2^, root mean squared error (RMSE), relative root mean squared error (RRMSE), the Pearson correlation coefficient (r), and the mean bias for model evaluation. RRMSE is the ratio of the root-mean-squared error to the mean of observed values, which normalizes RMSE by the target variable range and presents it as a percentage for easy cross-biome comparison. More accurate models are expected to have higher correlation coefficients and smaller RMSEs, RRMSEs and bias errors when compared with data. Analyses were conducted in the open-source environments R (version 4.1.0) and Python (version 3.8.10).

#### 2.2.6 Parameter sensitivity tests

For simplicity, in our main analysis, we fitted a single value for the parameter *σ* across all sites and years. However, we also investigated the variation of *σ* with environment (Table S5). Additionally, given the potential effects of satellite data uncertainty on *σ*, we fitted *σ* at flux sites based on MODIS and Copernicus data separately (Table S6). Based on the P model derived GPP and the fAPAR and LAI products of MODIS, GLOBMAP, and AVHRR, we further explored the spatial variation of σ on a global scale (Fig. S4) to assess its sensitivity to input GPP and satellite data.

### 3 Results

### 3.1 LAI variability across scales

The LAI model can capture the observed pattern of variability in LAI across temporal scales (Table 2, Fig.2). It explains 67 % of the variance in LAI with data aggregated to seven-day means (53863 data points) when driven by flux tower-derived GPP (Model_Flux_), and 60 % of variance when driven by the P model-derived GPP (Model_Prognostic_). Seasonal and annual variations are also well simulated (*R*^2^: 70% to 87 % for Model_Flux_ and 67% to 70% for Model_Prognostic_). Still more variance is explained for LAI across sites, with 90% for Model_Flux_ and 72% for Model_Prognostic_. Site-level interannual variations are less well simulated (*R*^2^: 43% for Model_Flux_, and 34% for Model_Prognostic_) (Fig.S5).

The LAI model also captures the seasonal and interannual variations of annual LAI for each biome (Figs 3,4, Table 3). The LAI simulated by Model_Flux_ (Fig. 3) agrees well with satellite-derived LAI, especially for MF (R^2^ = 58%; RRMSE = 18%), GRA (R^2^ = 69%; RRMSE = 27%), DBF (R^2^ = 79%; RRMSE = 22%), EBF (R^2^ = 56%; RRMSE = 23%) and WSA (R^2^ = 44%; RRMSE = 26%) and CSH (R^2^ = 72%; RRMSE = 15%). Observed LAI dynamics for ENF lag simulated LAI, with a delay of up to one month in spring (R^2^ = 39%; RRMSE = 15%). The ability of the model to simulate LAI in arid biomes (tropical savanna, open shrubland) were relatively poor; nonetheless, the model still accounted for 38% and 51 % of the variance in satellite-derived LAI, with RRMSE of 27% and 30% respectively, in these biomes. Compared to the LAI model driven by flux tower-derived GPP (Model_Flux_), the simulated LAI using P model-derived GPP (Model_Prognostic_) tends to overestimate LAI in early spring, especially for MF (R^2^ = 54%; RRMSE = 27%), GRA (R^2^ = 63%; RRMSE = 26%), ENF (R^2^ = 55%; RRMSE =2 4%) and DBF (R^2^ = 48%; RRMSE = 28%). This bias reflects the P model’s known tendency to overestimate GPP in the early part of the growing season, probably because it does not take account of the time required for new leaves to become fully functional under conditions of high light and low temperature (Stocker et al., 2020; Luo et al., 2023; Fig. S7). The underestimation of GPP (but not LAI) at some savanna sites (Figs 4e, S7) is likely because the sub-daily model does not consider C_4_ photosynthesis (Mengoli et al., 2022).

**Table 3.**
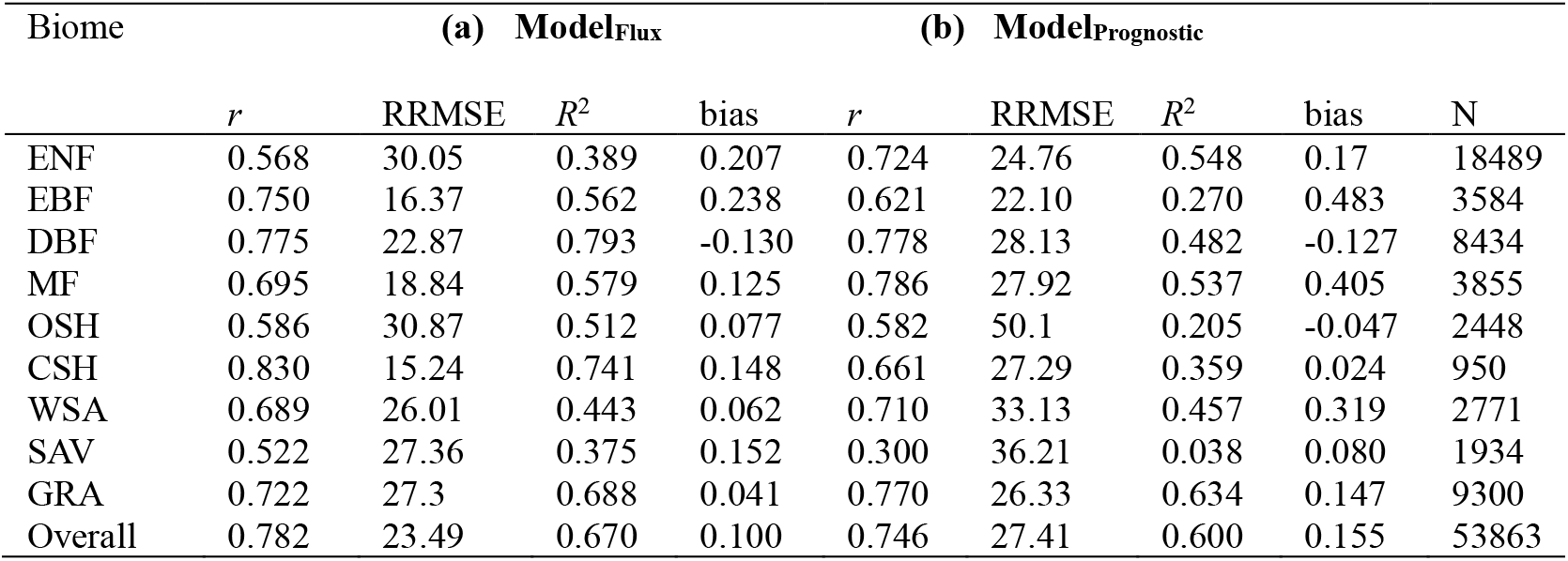
Observed versus modelled seven-day mean Leaf Area Index (LAI) by biome. **(a)** Model_Flux_: Inputs are flux-derived *A*_0_ and annual peak LAI obtained from the remote sensing data; **(b)** Model_Prognostic_: Inputs are P model-derived *A*_*0*_ and predicted annual peak LAI from Eqn 2. Reported metrics are the Pearson correlation coefficient (*r*), relative root means square error (RRMSE), *R*^2^ and bias. **CSH**, Closed shrubland; **DBF**, Deciduous broadleaf; **EBF**, Evergreen broadleaf; **ENF**, Evergreen needleaf; **GRA**, Grassland; **MF**, Mixed Forest; **OSH**, Open shrubland; **SAV**, Savanna; **WSA**, Woody savanna.

| Biome | (a) Model<sub>Flux</sub> |  |  |  | (b) Model<sub>Prognostic</sub> |  |  |  |  |
| --- | --- | --- | --- | --- | --- | --- | --- | --- | --- |
| | $r$ | RRMSE | $R^2$ | bias | $r$ | RRMSE | $R^2$ | bias | N |
| ENF | 0.568 | 30.05 | 0.389 | 0.207 | 0.724 | 24.76 | 0.548 | 0.17 | 18489 |
| EBF | 0.750 | 16.37 | 0.562 | 0.238 | 0.621 | 22.10 | 0.270 | 0.483 | 3584 |
| DBF | 0.775 | 22.87 | 0.793 | -0.130 | 0.778 | 28.13 | 0.482 | -0.127 | 8434 |
| MF | 0.695 | 18.84 | 0.579 | 0.125 | 0.786 | 27.92 | 0.537 | 0.405 | 3855 |
| OSH | 0.586 | 30.87 | 0.512 | 0.077 | 0.582 | 50.1 | 0.205 | -0.047 | 2448 |
| CSH | 0.830 | 15.24 | 0.741 | 0.148 | 0.661 | 27.29 | 0.359 | 0.024 | 950 |
| WSA | 0.689 | 26.01 | 0.443 | 0.062 | 0.710 | 33.13 | 0.457 | 0.319 | 2771 |
| SAV | 0.522 | 27.36 | 0.375 | 0.152 | 0.300 | 36.21 | 0.038 | 0.080 | 1934 |
| GRA | 0.722 | 27.3 | 0.688 | 0.041 | 0.770 | 26.33 | 0.634 | 0.147 | 9300 |
| Overall | 0.782 | 23.49 | 0.670 | 0.100 | 0.746 | 27.41 | 0.600 | 0.155 | 53863 |

### 3.2 Annual mean LAI at flux sites

Model_Prognostic_ tends to overestimate annual mean LAI in more arid biomes (Fig. 5b) compared to Model_Flux_ (Fig. 5a), likely due to higher predicted annual maximum LAI in SAV, GRA, WSA, and OSH (Figs 3, 4). Conversely, the underestimation of annual mean LAI for EBF and DBF is likely caused by a lower predicted annual maximum LAI and *A*_0_ (Figs 3, 4). The RMSE between satellite-derived and modelled LAI (Fig.5) is < 0.5 m^2^m^-2^ for 45% of the sites, < 1 m^2^m^-2^ for 90% of the sites, and < 1.5 m^2^m^-2^ at 98% of the sites. There are four sites where the RMSE exceeds 1.5 m^2^m^-2^, one SAV site (CG-Tch, RMSE = 1.69 m^2^m^-2^) and three EBF sites (AU-Tum: 2.19 m^2^m^-2^; BR-Sa3: 1.87 m^2^m^-2^; AU-Wac: 1.58 m^2^m^-2^).

### 3.3 Global-scale modelling

Our model captures MODIS-derived multi-year average LAI over 2001−2019 across most of global land surface covered (or partly covered) by natural vegetation. It somewhat underestimates LAI in Amazonia and northeastern Asian temperate deciduous forest, and overestimates LAI in arid regions including parts of southern Africa and South America (Figs 7, S6, S9). The simulated results are highly consistent with the latitudinal patterns shown in MODIS-derived products with multi-year mean LAI, with a correlation coefficient of 0.91 (Figs 7c-d). The seasonal time course of LAI on a global scale is also well simulated, apart from the “early spring effect” in the northern hemisphere, and a slight underestimation of global LAI in January

**Figure 7.**
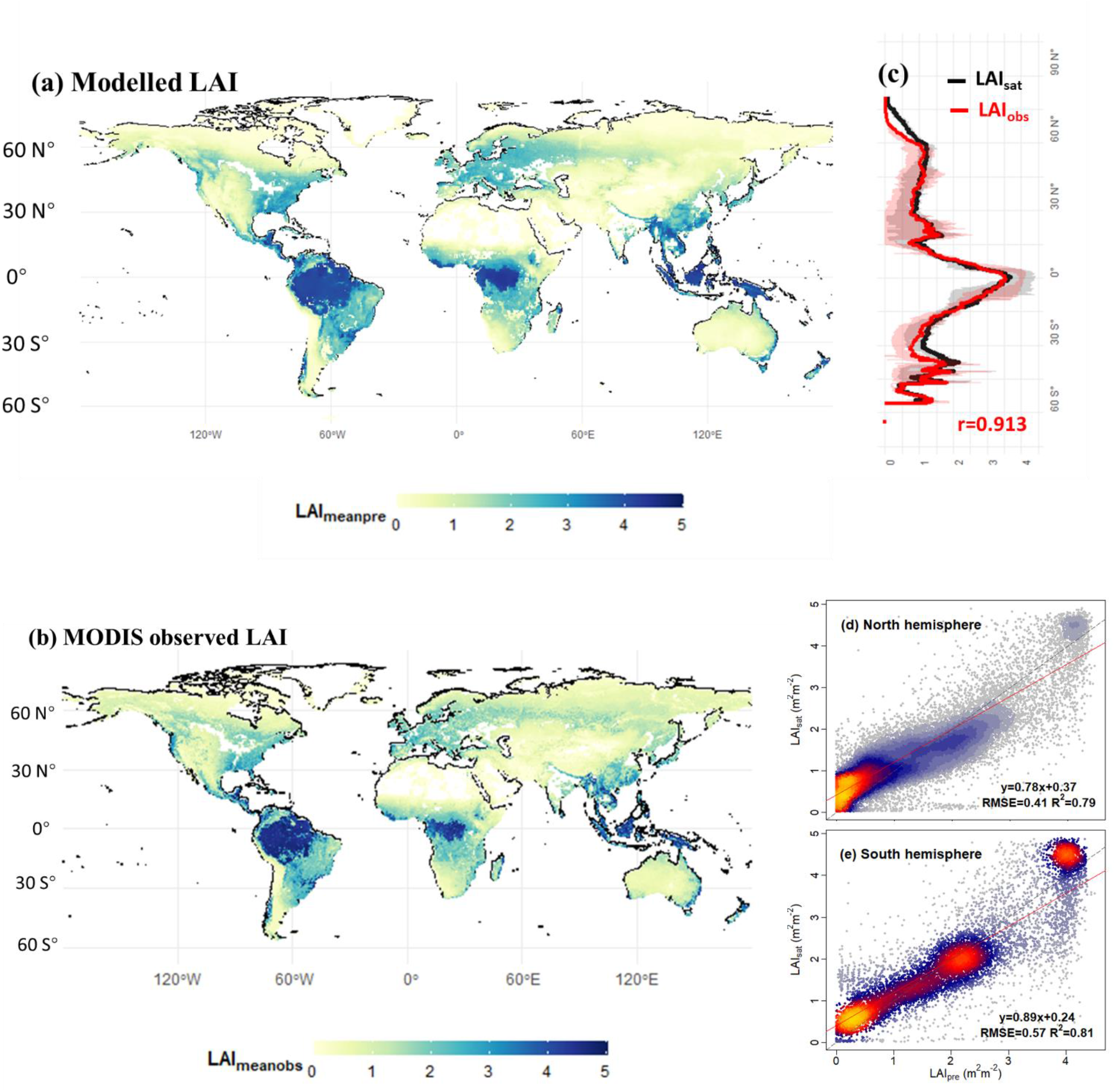
Spatial distributions of multi-year average LAI (2000 to 2019) from (a) our modelled results and (b) MODIS products, and (c) their latitudinal variation. Regressions are shown for the comparisons between our modelled results and MODIS products in the northern (d) and southern (e) hemispheres. The red and grey lines in (d)-(e) are the regression lines and 1:1 lines, respectively. Areas dominated by croplands snow/ice, and non-vegetated areas are shown in white in panels (a)-(b).

(Fig. S10). Our model is better than the 15 TRENDY models at predicting multi-year average LAI for 2001−2019 (Fig. S11), annual average LAI time series (Figs S12, S13, S14), and seasonal variations of LAI (Figs S15, S16, S17).

### 3.4 Effects of parameter variation

The predicted values of *m* captures the variation of observed values well (Slope = 0.90; R^2^ = 91%; RRMSE = 46%) for all biomes together (Fig. 8c). Separately fitted values of *σ* for EBF and ENF are higher (EBF: σ = 0.95; R^2^ = 96%; RRMSE = 15%; ENF: σ = 0.82; R^2^ = 88%; RRMSE = 37%) than those for DBF (σ = 0.67; R^2^ = 89%; RRMSE = 59%) and MF (σ = 0.63; R^2^ = 92%; RRMSE = 50%) (Fig. 8a; Table 3). CSH, GRA, OSH, SAV, and WSA show *σ* values in a relatively narrow range, from 0.69 to 0.82 (Fig. 8b). Even though *σ* values decrease with temperature and increase with PPFD, these variations are small (Table. S7). The σ values fitted using different satellite products at flux sites and global scales also differ very little (Tables S5, S6; Figs S4, S6). These results show that σ is a relatively robust parameter and can be used to derive a general expression for predicting the quantitative relationship between steady-state LAI and steady-state GPP (Eqn 10), applicable to all vegetation types.

**Figure 8.**
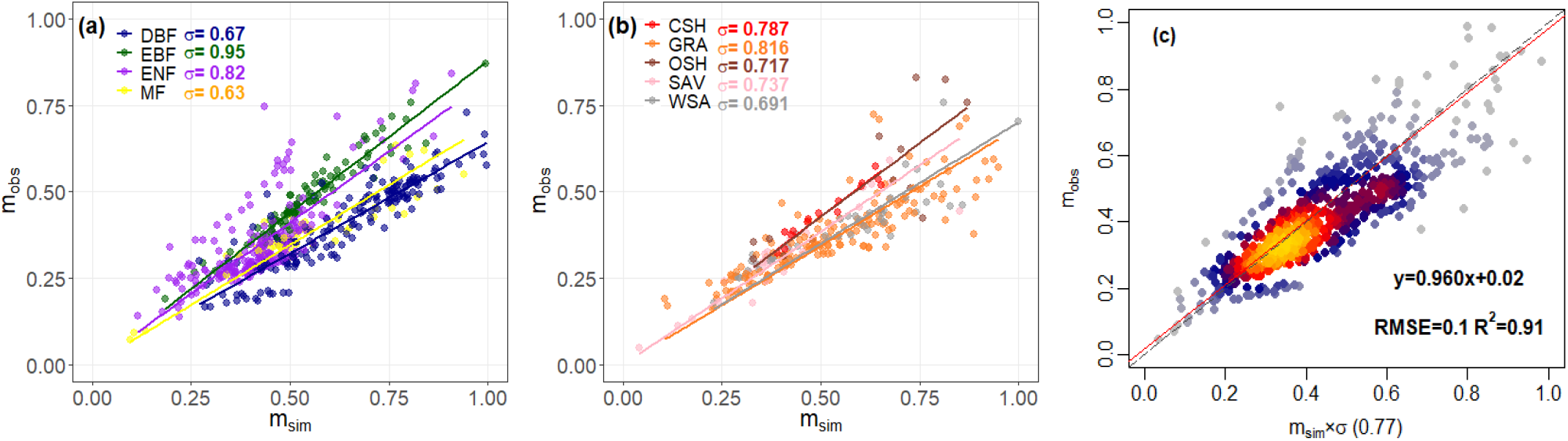
Variation of the parameter *m* for different biomes **(a)-(b)**, and all biomes together **(c). CSH**, Closed shrubland; **DBF**, Deciduous broadleaf; **EBF**, Evergreen broadleaf; **ENF**, Evergreen needleaf; **GRA**, Grassland; **MF**, Mixed Forest; **OSH**, Open shrubland; **SAV**, Savanna; **WSA**, Woody savanna.

## 4 Discussion

We have developed a universal global phenology model that predicts the daily time course of LAI from climate variables alone. The model is based on the theoretical approach introduced by Xin et al. (2018, 2020), which can be considered as an eco-evolutionary optimality hypothesis as it proposes that the dynamics of leaf display is closely coupled (with some time lag) to the time course of the productive capacity of leaves, which in turn is determined by light and weather conditions. We have combined this theoretical approach with a parsimonious and well-tested GPP model, the P model (Wang et al., 2017; Stocker et al., 2020; Mengoli et al., 2022), and a separately validated top-down model to predict the annual maximum LAI (Zhu et al., 2023; Cai et al., 2024). We have also developed a semi-empirical equation predicting the quantitative relationship between steady-state LAI and steady-state GPP, expressed as the parameter *m*, that is applicable across all biomes.

Our approach differs fundamentally from that of classical mechanistic phenological models, which focus on the triggers of phenological phase transitions (Chuine et al., 2000; Chuine and Régnière, 2017; Chuine, 2021). Mechanistic models can achieve high accuracy for phenological predictions at regional (Chuine et al., 2000) and species (Chuine et al., 2010) levels. Our optimality-based approach, however, may be better adapted for global vegetation and land-surface modelling.

Many studies have demonstrated a positive correlation between LAI and photosynthetic activity at various temporal and spatial scales (Wu et al., 2016; Chen et al., 2020; Bian et al., 2023). Our results provide further support for the hypothesis of interdependence between the time course of LAI and the photosynthetic potential of leaves. This hypothesis provides a robust way to predict seasonal leaf allocation from the conditions supporting photosynthetic productivity, instead of trying to predict the timing of phenological events (McMaster and Wilhelm, 1997; Badeck et al., 2004; Cleland et al., 2007). The resulting model is indifferent to the specific mechanisms used by plants in different environments to achieve optimality. The model also avoids explicitly simulating the dynamics of C allocation to leaf growth. One recent study found that leaf C accumulation accelerates in the early green-up phase (due to both increased photosynthesis and a greater fraction of GPP allocated to leaves) and declines in the late green-up phase (Meng et al., 2023). However, the theory of dynamic C allocation is incomplete (Hartmann et al., 2020) and there is insufficient information that would allow a time-varying allocation fraction to be modelled globally. Relying on the interdependence of LAI time series and photosynthetic potential provides an effective alternative, as shown by Zhou et al. (2023), and in the present study.

Our LAI simulation framework tends to overestimate LAI values in spring in cold-climate regions, due to a known deficiency of the P model (Stocker et al., 2020; Mengoli et al., 2022), which may be related to the fact that full photosynthetic activity lags behind leaf development in these environments (Croft et al., 2015; Luo et al., 2023). The model as presented here has benefited from including a climatically variable *f*_0_ parameter (the fraction of antecedent precipitation hst is used by plants), compared to the original formulation for the calculation of seasonal maximum fAPAR (Zhu et al., 2023; Cai et al., 2024). However, the model stiill shows some tendency to overestimate LAI in arid regions.

## 5. Conclusions

We have developed a prognostic and globally applicable phenology model which can predict daily LAI based on climate variables alone. This model is based on an assumed linear relationship between “steady-state” values of LAI and GPP. The exponential moving average method is used to account for the time lag between leaf allocation and steady-state LAI. The model captures the spatiotemporal patterns of satellite-derived LAI across biomes, both at eddy-covariance flux measurement sites, and in terms of global patterns. Some problems remain: the model predicts too early spring green-up in some cold-winter climates and tends to overestimate LAI in dry climates. Nonetheless, the model performs better than the 15 DGVMs participating in the TRENDY project. Applications of the P model to date have used satellite-derived green vegetation cover indices as input. The model presented here allows us to predict the seasonal time course of both LAI and GPP across all natural ecosystems independently of satellite data − representing an important step towards the development of a “first principles” next-generation land surface model that will be both simpler, and more accurate, than those currently in use.

## Code availability

The P-model is implemented as a Python package (pyrealm) and available through GitHub (https://github.com/ImperialCollegeLondon/pyrealm/tree/develop/.github). Results shown here correspond to pyrealm version 0.9.0 and the subdaily module of the P model.

## Data availability

The flux tower dataset can be accessed from the FLUXNET 2015 Tier Two datasets and ONEFLUX dataset (http://fluxnet.fluxdata.org/, last access: 15th July 2023) (Pastorello et al., 2020). MODIS LAI and MODIS fAPAR data are from the MODIS MOD15A2H Leaf Area Index/FPAR product (Jiang et al., 2016). Copernicus LAI and fAPAR data were derived from Copernicus Global Land Products (https://gbov.acri.fr, last access: 20^th^ July 2023).

## Author contributions

ICP, SPH and WH designed the research; BZ and ICP developed and refined the methodology; BZ, ZZ, and WC carried out data processing and analyses; all authors contributed to the interpretation of the results. BZ created the graphics under the supervision of ICP and wrote the first draft. All authors contributed to the final version.

## Acknowledgements

This research is a contribution to the Land Ecosystem Models based On New Theory, obseRvations and ExperimEnts (LEMONTREE) project which received support through Schmidt Sciences, LLC. BZ is funded by LEMONTREE. WC and ICP acknowledge support from the European Research Council under the European Union’s Horizon 2020 research and innovation programme (Grant Agreement No: 787203 REALM). ZZ acknowledges support from the National Natural Science Foundation of China (32022052, 91837312, 31971495) and the Tsinghua University Initiative Scientific Research Program (20223080041). Many thanks to Dr David Sandoval for his suggestions and helps in improving the annual maximum LAI simulation in arid regions. Also many thanks to Dr Giulia Mengoli for help with the application of the subdaily GPP model.

